# CRI-SPA – a mating based CRISPR-Cas9 assisted method for high-throughput genetic modification of yeast strain libraries

**DOI:** 10.1101/2022.07.19.500587

**Authors:** Helén Olsson, Paul Cachera, Hilde Coumou, Mads L. Jensen, Benjamín J. Sánchez, Tomas Strucko, Marcel van den Broek, Jean-Marc Daran, Michael K. Jensen, Nikolaus Sonnenschein, Michael Lisby, Uffe H. Mortensen

## Abstract

Biological functions are orchestrated by intricate networks of interacting genetic elements. Predicting the interaction landscape remains a challenge for systems biology and the identification of phenotypic maximas would be of great benefit to synthetic biology. Thus, new research tools allowing simple and rapid mapping of sequence to function are required to forward these research fields. Here, we describe CRI-SPA, a method allowing the transfer of a chromosomal genetic feature from a donor strain to arrayed strains in large libraries of *Saccharomyces cerevisiae*. CRI-SPA is based on mating, <u>CRI</u>SPR-Cas9-induced gene conversion and <u>S</u>elective <u>P</u>loidy <u>A</u>blation and is executed within a week. We demonstrate the power of CRI-SPA by transferring four genes responsible for the production of betaxanthin, a yellow biosensor for the morphine precursor L-DOPA, into each strain of the yeast knock-out collection (≈4800 strains), providing a genome-wide overview of the genetic requirements for betaxanthin production. CRI-SPA is fast, highly reproducible, can be massively parallelized with automation and does not require selection for the transferred genetic feature.

## Introduction

Baker’s yeast *Saccharomyces cerevisiae* is an important biological model organism and a key production host in industrial biotechnology. *S. cerevisiae* was the first eukaryote to be fully sequenced (Goffeau et al., 1996; Cherry et al., 1997) and its simple lifecycle as a single cell organism, with a highly developed genetic toolbox have positioned this yeast as a frontrunner in the field of systems biology. However, the unpredictability of phenotypic effects resulting from combinations of different genetic traits still challenges our fundamental understanding of yeast and its engineering. The introduction of CRISPR for yeast engineering (DiCarlo et al., 2013) promises to relieve this bottleneck by accelerating the systematic construction and testing of genetic variants. For example, in numerous genome-wide engineering projects, CRISPR-based strategies have been developed to create and screen libraries of strains (Replogle et al., 2020; Lian et al., 2019; Sharon et al., 2018; Bao et al., 2018; Han et al., 2017; Wong et al., 2016; Garst et al., 2017; Sadhu et al., 2018). These screens generally follow a one-pot workflow where the library is pooled, edited and challenged before being resolved by FACS and next generation sequencing (Bock et al., 2022; Shalem et al., 2015). However, library pooling is prone to bias as it selects for fast growth and traits of interest might be lost if they are accompanied by a reduction in growth fitness. Similarly, when recovering the best candidates from a one-pot screen, the top pool might be saturated by a few strongest variants, which limits the full network of interactions to be uncovered (Savitskaya et al., 2019). A more useful output would be achieved if the members of the library could be assessed individually to produce the full overview of the genetic effects impacting the system.

Synthetic Genetic Array (SGA) queries strains individually in a process based on arrayed mating, meiosis, sporulation and double marker selection for the desired gene combination. This method has been particularly valuable for the identification of genetic interactions among all double and a selected number of triple gene knock-outs (Kuzmin et al., 2021; Costanzo et al., 2016). However, the reliance of SGA on meiosis, the longest step in SGA (4-7 days), and the use of selectable markers might become inadequate for multiple applications. For example, correct chromosome segregation in the first meiotic division requires high levels of homologous recombination (Lichten, 2001), which may lead to undesirable genomic rearrangements. Flawed meiosis is also frequently known to generate aneuploids (Adames et al., 2019; Jaffe et al., 2017), which could compromise further analyses. In addition, selectable markers may influence expression of neighboring genes and become a limitation for multiplexing. A faster method, which is independent of meiosis and does not rely on marker selection, would therefore be desirable.

We have developed a new screening platform that allows a genetic trait to be queried in large arrayed yeast libraries. Our method (Figure 1), CRI-SPA, combines <u>CRI</u>SPR (Jinek et al., 2012; DiCarlo et al., 2013) with selective ploidy ablation, <u>SPA</u> (Reid et al., 2011). In CRI-SPA, a (marker-free) genetic feature of interest is efficiently transferred from a <u>C</u>RI-SPA <u>D</u>onor <u>S</u>train (CDS), to the strains in a library in a process involving mating, Cas9 induced gene conversion and haploidization by SPA. CRI-SPA can be massively parallelized using a pining robot and executed in a week with as little as 4 hours handling time. Since it relies on image analysis for data extraction, the main cost associated with CRI-SPA comes from media and plastic consumables.

**Figure 1.**
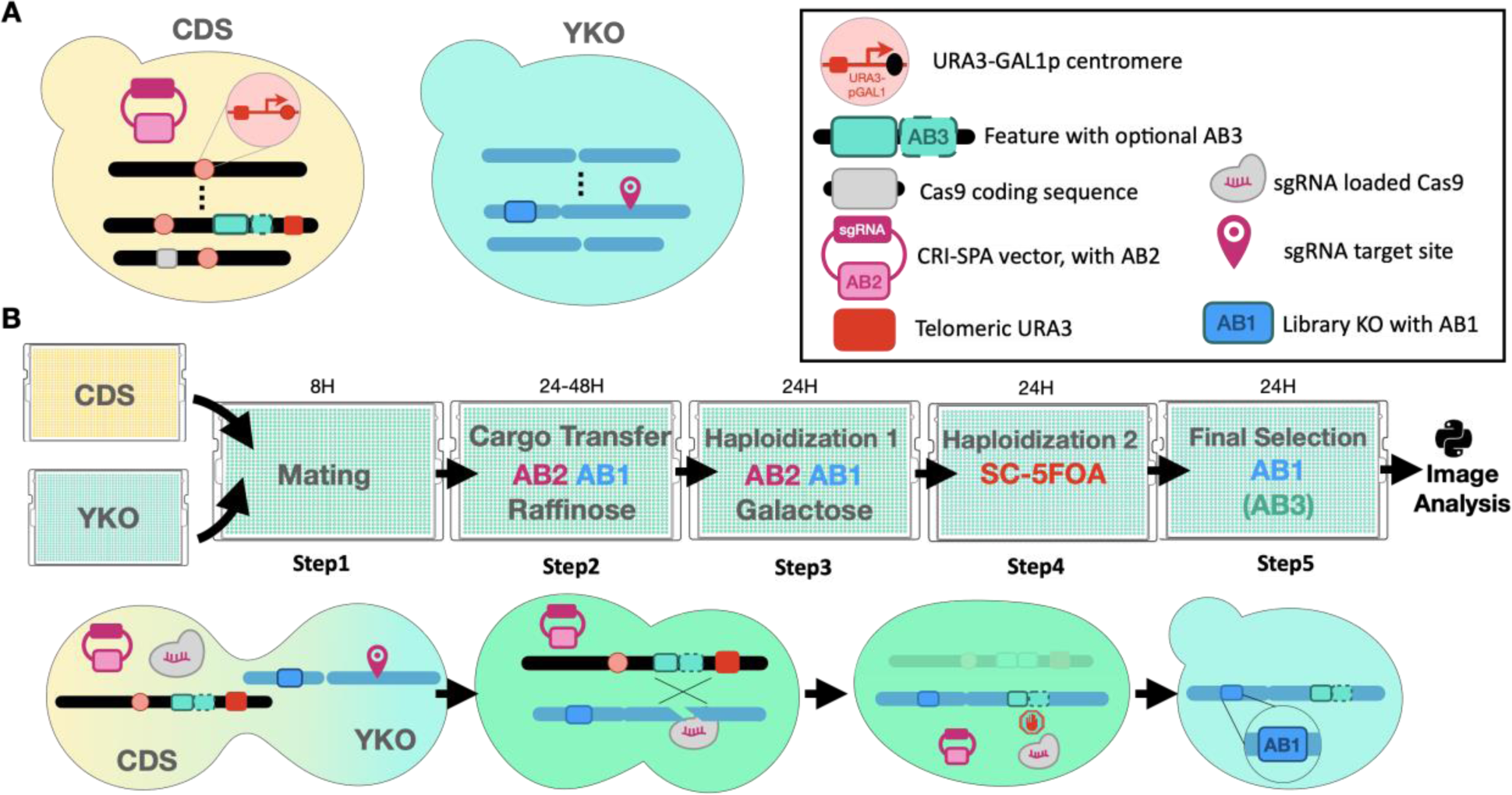
The CRI-SPA gene-transfer system. A) Left: The CRI-SPA donor strain (CDS). The CDS contains: a genetic feature of interest coupled to an antibiotics marker, AB3 (optional), a *URA3* gene between the genetic feature of interest and the telomere, *URA3-GAL1p* cassettes at the centromeres of all chromosomes, a *cas9* and a selectable CRI-SPA plasmid maintained by another antibiotics marker (AB2) and encoding a sgRNA targeting the insertion site of the genetic feature (blocked in the CDS). Center: The library strains (here the YKO) contain the antibiotic marker 1 (AB1) and a target site for the Cas9-sgRNA CRISPR nuclease. Graphics of the individual genetic components are shown in the legend box. (B) The individual steps of the CRI-SPA procedure (see main text for details). **Step 1**, a CDS strain is pinned onto all strains of the library plates and incubated for mating. **Step 2**, diploids are exhausted for glucose via growth on raffinose and selected for the marker in the library strains (AB1) and the marker on the CRI-SPA plasmid (AB2). At this stage, the target site in the recipient strain is cleaved by Cas9-sgRNA. Repair of the resulting DNA DSB by gene conversion using the corresponding donor site as template transfers the genetic feature to the recipient locus. Note that the target site of the Cas9-sgRNA is destroyed by the genetic feature of interest. **Step 3**, galactose induced SPA eliminates the donor chromosomes. **Step 4,** haploid cells containing only recipient chromosomes are selected on 5- FOA. **Step 5,** the marker of the recipient (AB1) and the genetic feature (AB3 optional) are selected yielding the final library.

Here, we use the method to transfer four genes responsible for the synthesis of the yellow plant metabolite betaxanthin, a biosensor for the morphine precursor L-DOPA (DeLoache et al., 2015), into the 4800 strains of the yeast knock-out collection (Giaever et al., 2002). In doing so, we unveil the full betaxanthin pathway-host gene interaction landscape. Most prominently, we demonstrate that mutations impairing mitochondrial functions consistently produce strains that are significantly more yellow than the reference strain.

## Results

### Experimental steps of the CRI-SPA method

Prior to CRI-SPA, a CDS is constructed from our Universal CRI-SPA strains (UCSs), (supplementary Figure 1). Once the CDS is constructed (Supplementary Materials and Methods.), the CRI-SPA procedure is performed in five steps during 6 days (Figure 1). In **step 1,** donor and recipient strains mate overnight on YPD medium. In **step 2**, strains are transferred onto raffinose medium for 48 hours. At this stage, the Cas9- sgRNA induced double-strand break at the recipient locus is repaired through HR using the donor chromosome as a repair template. This results in the transfer of the genetic feature from the donor to the recipient chromosome. In **step 3,** transition from raffinose to galactose allows for the sudden induction of the *GAL1* promoter which disrupts the centromeres of all donor strain chromosomes resulting in their selective loss (Reid et al., 2011). In **step 4**, cells that have lost all donor chromosomes are selected by transferring the strains to solid SC medium containing 5- FOA. This medium also counterselects undesired recombinant strains resulting from DNA DSB repair involving crossing-over or break induced replication at the target locus (Supplementary Figure 2). As a result, step 4 generates haploid strains that solely contain the recipient strain’s chromosomes as well as the genetic feature of interest. In the final **step 5,** recipient cells are selected e.g. by the *kanMX* marker of the YKO library; and optionally, the genetic feature of interest is also selected for if it is accompanied by a marker. Conveniently, we note that CRI-SPA can be executed by hand or scaled with a pinning robot to accommodate a range of budgets and throughputs.

### Transfer of *ade2*Δ*::hphNT1* into a selected set of arrayed gene deletion mutants

The present CRI-SPA procedure (Figure 1) is the result of several rounds of optimization. In this process, transfer of an *ade2*Δ*::hphNT1* cassette from the donor strain CDS-*ade2*Δ to a subset of the YKO library was employed as a test screen. Advantageously, *ade2Δ* strains produce an easy-to-score red phenotype (Wintersberger et al., 1995; Smirnov et al., 1967), which may be epistatically blocked by mutations upstream in the purine pathway, e.g. *ade3* (Roman, 1956). Hence, successful interchromosomal transfer of *ade2*Δ*::hphNT1* into the YKO library should produce mostly red colonies, but should also identify epistatic genetic interactions, like *ade3*Δ, which should appear as white colonies. In addition, production of *ade2* cells provide an excellent test-bed to validate the robustness of CRI-SPA as they accumulate toxic intermediates causing a fitness burden (Chaudhuri et al., 1997). Hence, since CRI-SPA colonies are composed by a population of cells surviving the procedure, the efficiency of the method can be assessed as even a few undesired *ADE2* escapers in a colony may penetrate in the end of the screen due to their competitive edge.

Successful CRI-SPA mediated allele transfer requires efficient production of DNA DSBs at the target locus by the CRISPR nuclease. Prior to the *ade2*Δ::hphNT1 transfer experiment, we therefore demonstrated that the *ADE2* CRI-SPA vector (pHO-*ADE2*) produced efficient CRISPR nuclease activity to efficiently cleave the *ADE2* locus as judged by the Technique to Assess Protospacer Efficiency (TAPE) experiment (Vanegas et al., 2017, supplementary Figure S3). In parallel with this test, we used the same vector to mediate integration of the *ade2Δ:*:hphNT1 cassette into an UCS1 MATα strain to produce CDS-*ade2*Δ.

Next, CDS-*ade2*Δ was used in a CRI-SPA experiment screening the strains on plate 9 of the YKO library (*MAT<u>a</u>*), which contains 376 gene deletions, including *ade3*Δ::kanMX, a negative interaction control. As a starting point, we first used the standard SPA protocol (Reid et al., 2011), but with an additional step to select for diploids (Supplementary Figure S4A). After CRI-SPA, several solid red colonies were observed, but many were still white or pink, (Supplementary Figure S4B). Note that CRI-SPA produces colonies containing a population of cells rather than clonal strains. We therefore restreaked pink colonies on solid SC medium which yielded both white and red cells indicating the presence of clones with the expected *ade2Δ* mutation as well as white escapers. Importantly, we noticed that early in the CRI-SPA procedure, almost all colonies were red. This hinted that rare white cells later outcompeted red cells in a colony. We therefore repeated the experiment using plates with increased adenine levels to examine whether downregulation of the purine pathway would improve CRI-SPA efficiency. This modification of the protocol decreased, but did not eliminate, the presence of white and pink colonies (Olsson, 2020). Next, we examined whether these colonies were due to escapers simply representing diploid cells that somehow passed through the 5-FOA selection procedure. However, this appeared not to be the case as the majority of streak purified white colonies failed to grow on SC-Ura plates (Olsson, 2020). Moreover, diagnostic PCR reactions of these colonies demonstrated that they contained both *ade2Δ*::hphNT1 and *ADE2* alleles indicating that they harbored two copies of chromosome XV. It has previously been shown that the SPA procedure may trigger endoreduplication (Reid et al., 2008) and it may therefore also occur during a CRI- SPA experiment. Indeed, flow cytometry analyses of white strains demonstrated that they appear to contain a population of 2N and 4N cells rather than 1N and 2N cells expected for a haploid strain (Supplementary Figure S5). Based on these results, we speculated that undesired white cells were most likely formed by CRI-SPA gene transfer taking place during or after endoreduplication in a manner where only one of the duplicated recipient chromosome XV copies were modified by gene conversion (Supplementary Figure S6A-C). Alternatively, we note that many strains in the gene deletion library suffer from aneuploidy (Hughes et al., 2000), which may also provide a source for recipient strains containing both *ADE2* and *ade2*Δ::hphNT alleles using a similar scenario for gene transfer (Supplementary Figure S6D). To address these possibilities in more detail, we examined three strain backgrounds, *hse1*Δ, *scp160*Δ, and *atp20*Δ, which all produced red and white colonies after streak purification, by full genome sequencing. As expected, the three red colonies contained the *ade2*Δ::hphNT allele, but not the CDS donor chromosome; and the three white strains contained two s288C based copies of chromosome XV, one containing the *ADE2* allele and one the *ade2*Δ::hphNT allele. Interestingly, gene transfer was accompanied by transfer of SNPs from W303 to s288C around the CRISPR induced DNA DSB site. In some cases, the SNPs were only found close to the DNA DSB site, but for others they appeared up to 20 kb away from the break site. Hence, DNA DSB repair in this CRI-SPA experiment appears to be accompanied by short and long gene-conversion tracks indicating that repair may occur by different pathways e.g. synthesis-dependent strand-annealing (short tracks) and break-induced-replication pathways (long tracks; Yim et al., 2014). Lastly, we note that none of the other recipient chromosomes contained any W303 SNPs.

With these insights, we envisioned that background could be reduced by including a growth step on non-reducing/non-repressing raffinose to allow more time for CRISPR mediated gene conversion and at the same time set the stage for more efficient galactose induction (Johnston et al., 1994). With this protocol, we repeated the *ade2*Δ::hphNT*1* CRI-SPA transfer to the same subset of the yeast deletion library. This time we upscaled the plate format from 384 (YKO library format) to 1,536, to perform the transfer in 2x2 quadruplicates. After CRI-SPA, virtually all colonies were red. For about six genes (out of 376) we observed that one of the quadruplicates was conspicuously pink. These incomplete CRI-SPA events are rare, indicating that production of background Ade^+^ cells had been almost eradicated (Figure 2A). Importantly, and, as expected, the *ade2*Δ::hphNT1 *ade3*Δ*::kanMX* double mutant produced four white colonies (Figure 2A). Two additional white quadruplets were observed, representing *ade2Δ::*hphNT1 *met6*Δ*::kanMX* and *ade2*Δ::hphNT1 *scp160*Δ::kanMX double mutants. The fact that none of the two double mutations were expected to cause a white phenotype suggested that these results were artifacts. Indeed, restreaking of these white colonies demonstrated that they contained cells that were able to form red colonies when streaked out into single colonies on solid selective media (G418 and hygromycin). Lastly, we observed a few mutant strains that did not produce any growth after CRI-SPA. Such mutants may represent strains that fail to go through CRI-SPA, e.g. if they fail to mate; or if they are synthetic lethal in combination with *ade2Δ*::hphNT1.

**Figure 2.**
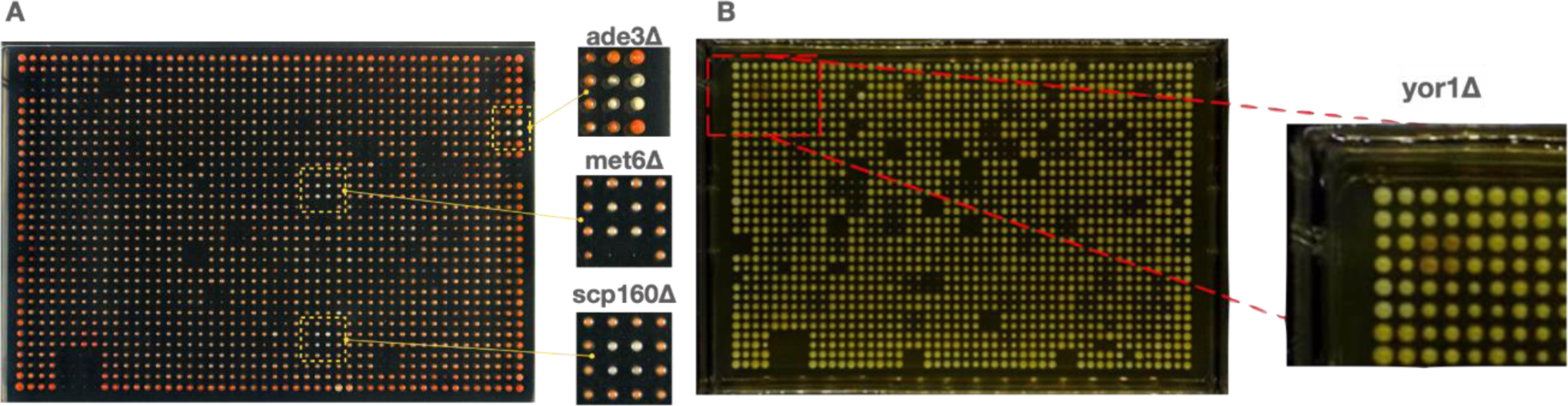
Pilot CRI-SPA experiments. **(A)**Transfer of an *ade2::hphNT1* mutation into a selected plate 9 of the YKO library (pinned in quadruplicates). Two quadruplicates that appear white (*ade2*Δ*::kanMX* and *scp160*Δ*::kanMX*, insets) are highlighted in yellow boxes and shown in amplification to the right as indicated. **(B)** Transfer of *CYP76AD1* and *DOD* for betaxanthin production into the full YKO library. 4 of 13 plates are shown. The section of plate 8 (quadruplicate of 386 format) containing *yor1*Δ*::kanMX* (position B2) is marked with a red box and shown in amplification to the right.

### Introduction of the betaxanthin pathway into a genome-wide library of gene deletion strains

We next decided to develop a setup allowing transfer of a multi-gene cassette harbored within the well-defined XII-5 integration site (Mikkelsen et al., 2012) of a CDS to a recipient strain library by CRI-SPA. For this purpose, we constructed Universal CRI-SPA type 2 strains (UCS2). In addition to the 16 centromere-adjacent *KlURA3* makers, the UCS2 harbors an additional *Kl_URA3* marker between site XII-5 and the telomere (see M and M). This design allows for the counter selection of clones resulting from Cas9 induced gene conversion at XII-5, which is accompanied by undesirable crossing over (Supplementary Figure S2). To set the stage for CRI-SPA transfer at XII- 5, we constructed a hphNT1 marker-based CRI-SPA plasmid (pHO-XII-5) encoding an sgRNA that efficiently mediates cleavage at this site.

To demonstrate the usefulness of the CRI-SPA in cell factory development, we decided to map the genetic requirements, at single gene resolution, for the production of betaxanthin. This yellow and fluorescent plant secondary metabolite, can be produced in yeast via expression of two heterologous genes *CYP76AD1* and *DOD* (DeLoache et al., 2015). In a new pilot CRI-SPA experiment, *CYP76AD1*, *DOD* and the selectable natMX marker were inserted into XII-5 of UCS2 type α as one large transferable multi-gene cassette yielding CDS-btx1. We then used CDS-btx1 to transfer the betaxanthin expression cassette to the 4787 unique deletion strains of the YKO library using CRI-SPA. The experiment was performed in quadruplicate for a total volume of 19,148 transfer reactions and production of betaxanthin was estimated using the degree of yellow color of the colonies as a proxy. Encouragingly, the vast majority of colonies appeared yellow after CRI-SPA indicating that the CRI-SPA mediated transfer of the betaxanthin pathway into the library was successful (Figure 2B). Moreover, for most mutants (83.8%), all four quadruplicates produced viable colonies. We noted that colonies were subject to positional artifacts which are known to affect high density screens (Baryshnikova et al., 2010; Collins et al., 2006). Nevertheless, when ranking colony yellowness using a commercial image analysis software for measuring color intensity (see M & M), we identified a strain containing a deletion of *YOR1*, as the third top hit (Figure 2B). *YOR1* has previously been identified as an exporter whose deletion increased intracellular betaxanthin concentration (Savitskaya et al., 2019). This result confirms the ability of CRI-SPA to identify genes beneficial to the synthesis of betaxanthin.

Next, we made several adjustments to our screening protocol to reduce positional artifacts and to make the screen more amenable to automatic data extraction by image analysis. Data processing has previously been developed to correct for positional effects affecting colony size (Baryshnikova et al., 2010). However, such a pipeline remains to be adapted to colony color. Since we screened the library in quadruplicates, we reasoned that a simpler approach would be to average out positional artifacts by randomizing the positions of quadruplicates. Specifically, we shuffled the four replicates across the screen plates when upscaling the 384 arrays of the YKO library to a 1,536 format (Supplementary Figure S7).

To increase the dynamic range of our screen, we used a new CDS strain (CDS-btx2), which in addition to *CYP76AD1*, *DOD* and *natMX*, also contains *ARO4^K229L^* and *ARO7^G141S^* in the transferable multi-gene cassette. These two genes contain dominant mutations alleviating the negative tyrosine-feedback regulation on Aro4 and Aro7 (DeLoache et al., 2015; Luttik et al., 2008) ensuring an optimized metabolite flux towards betaxanthin. This effect should enhance the range of yellow color displayed by the colonies (Savitskaya et al., 2019). Hence, in this CRI-SPA experiment, we attempted the simultaneous transfer of five genes into the full YKO library in quadruplicate. As in the previous experiment, the vast majority, 88.6%, of the deletion strains produced three or more colonies after CRI-SPA. We then adapted an image analysis pipeline (Kamrad et al., 2020) to quantify the degree of yellow color intensity of each colony (Supplementary Methods) and rank all knock-outs that produced at least three colonies (Figure 3). This analysis demonstrated a strong signal to noise ratio (the Z-scores of the 16 top and bottom hits was 40.83, -6.93; respectively, Figure 3 - middle). The color difference between top, middle and bottom colonies were also strikingly different on the screen plates (Figure 3 - left). Since CRI-SPA colonies are not monoclonal, we isolated three candidates representing three different color levels and restreaked them on solid YPD medium with appropriate selection. Again, color differences between the sets were easily distinguished by photography, validating the position of the mutant strains in the color ranking scheme (Figure 3 - right).

**Figure 3.**
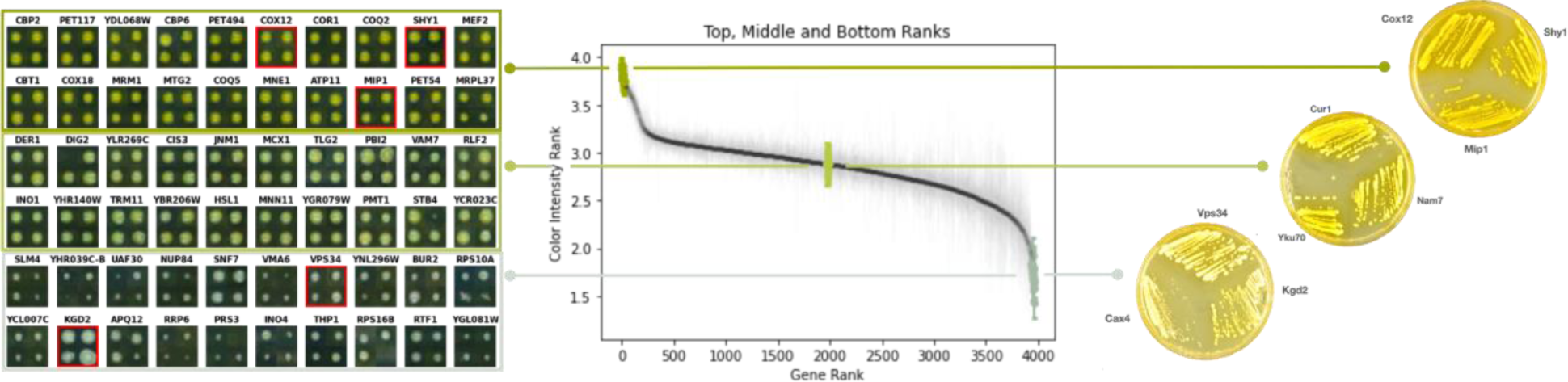
Genome-wide ranking of mutants according to yellow color intensity. **(Left)** Mosaics of selected mutants shown as quadruplicates from regions in the yellow color intensity ranking order **(Middle)** representing high, middle, or low degrees of yellowness as indicated. (**Right**) Mutants in red frames (left) were streak purified and clones were restreaked on solid YPD medium.

Confident that the screen readout was accurate for a handful of strains, we repeated the entire CRI-SPA experiment to validate its reproducibility. After CRI-SPA, the yellowness of the individual mutants obtained in this experiment, trial 2, were then plotted against the yellowness obtained for the mutants in the first experiment, trial 1. The clear linear correlation between the two trials (PCC= 0.765) demonstrates the reproducibility of CRI-SPA (Figure 4A). Moreover, we found that, from all successfully labeled genes in the two screens, 86 % (hypergeometric test p=1.09e-220) and 52 % (hypergeometric test p= 9.97e-83) of the 192-top and 192-bottom hits were common in the two trials, respectively. Hence, and especially for the top-hits, reproducibility was high.

**Figure 4.**
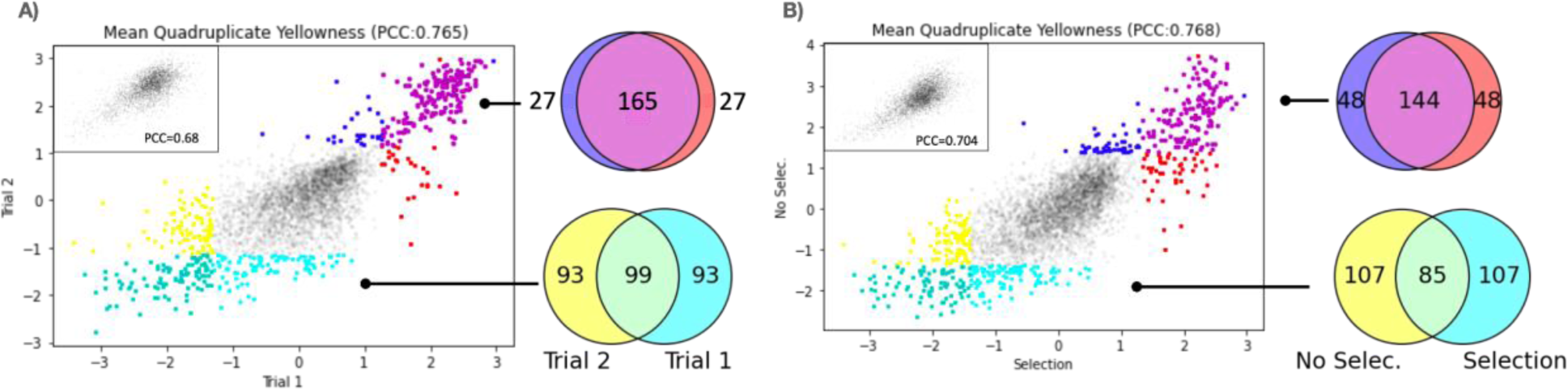
Reproducibility of betaxanthin CRI-SPA experiments in the presence and absence of pathway transfer selection. **(A)** The mean yellow color intensities obtained for each mutant in repeat 1 and repeat 2, which both were performed in the presence of natMX to select for the betaxanthin cassette genes. **(B)** The mean yellow color intensities obtained for each mutant in trial 1 and trial 3. Trial 3 was performed in the absence of natMX. The top and bottom 192 genes and their overlap across screens are coloured and displayed in flanking Venn diagrams. Insets show the correlation for colony fitness across screens.

In a third experiment, trial 3, we explored the possibility of transferring the five-gene betaxanthin cassette from the CDS to the full YKO library without selecting for the *natMX* marker, which is present in the cassette. In this screen, the CRI-SPA plasmid was maintained to use CRISPR restriction of the target site as the sole selection for genetic transfer. With this set-up, 95.1% of mutants produced at least 3 colonies after CRI-SPA mediated transfer of the betaxanthin cassette. Encouragingly, the yellowness score in this trial had a clear linear correlation with trial 1 (PCC=0.768, Figure 4B). Moreover, when the 192 most yellow and 192 least yellow mutants obtained in the two trials were compared, the top and bottom groups had 75% (hypergeometric test p=24.28e-181) and 44% (hypergeometric test p= 3.22e-70) overlap across screens, respectively (Figure 4B). Hence, the above analyses demonstrate that CRI-SPA is reproducible and that the combined effect of a mutation and a genetic feature can be reproduced even without selecting for this feature.

### Systems perspective of betaxanthin production

In contrast to one-pot screens, CRI-SPA produces a read-out for each individual strain within a library. We investigated whether this systematic gene labeling could reveal new system level patterns in betaxanthin production. We added the WT background strain (BY4741 encoding KanMX resistance at XI-3 insertion site) to the library and repeated the screen with antibiotic selection a third time, trial 4. We pooled the data of the three screens performed in the presence of selection obtaining a median of 11 replicates (12 is maximum) per gene. We first plotted yellowness of all strains against colony size using the latter feature as a proxy for fitness. We grouped genes according to their yellow score: the “yellow” group was defined as genes with the highest level of betaxanthin production (1.2 standard deviation greater than the screen mean, yellow points in Fig. 5). The “white” group was defined as genes with the lowest level of betaxanthin production (1.2 standard deviation below the screen mean, white points in Fig. 5). We observed that most members of the yellow group display a strong fitness penalty. We therefore defined a third group of mutants, the “cyan group” (Figure 5). This group contains many strains with little or no fitness penalty. Next, we examined whether specific cellular functions could be linked to betaxanthin levels by running gene ontology enrichment analyses (Klopfenstein et al., 2018) within the three groups. The cyan group did not produce any enrichment. This may not be surprising as the hits in this group were the least reproducible (Supplementary Figure S8). The white group, despite its broad fitness range, was enriched for terms describing the degradation machinery, translation, and vacuolar mechanisms. Most strikingly, the yellow group displayed a dramatic enrichment for genes associated with mitochondrial functions, particularly the respiratory chain, which might explain the fitness penalty observed in this group. Hence, this analysis demonstrates that CRI-SPA is able to identify a novel network of genes, which impact functions of a specific organelle in the cell, and which, when deleted, lead to reduced fitness and high betaxanthin levels.

**Figure 5.**
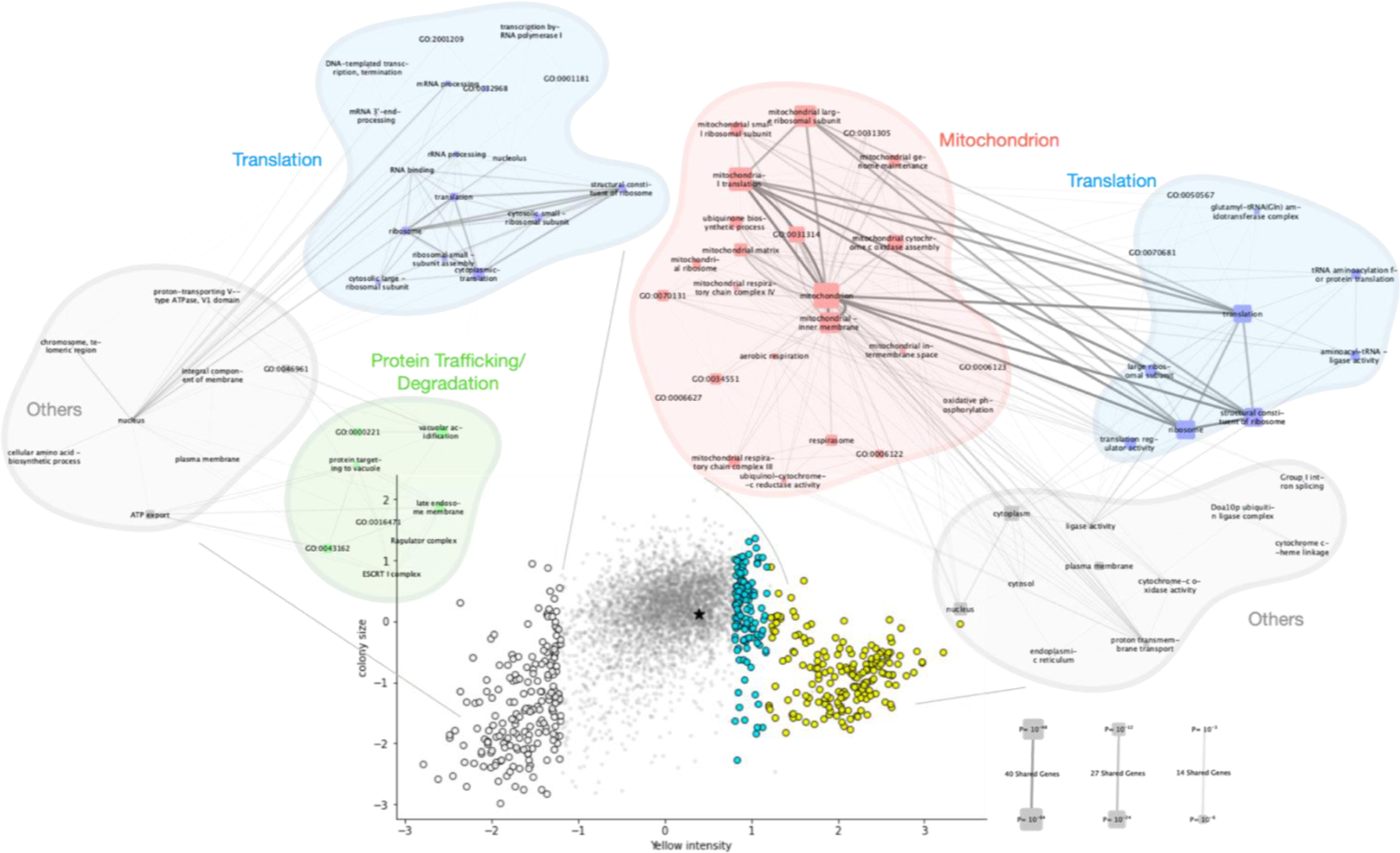
Betaxanthin yield and fitness relationship reveals mechanistic patterns in betaxanthin production. Colony size (i.e. fitness) vs Yellow Color intensity (i.e. Betaxanthin Yield). The results of Gene Ontology Enrichment Analyses on several subgroups of genes are shown as graphes for the bottom hits (white, 1.2 std below the screen mean) and top hits (yellow, 1.2 std above the screen mean). Node size indicates the significance of GO terms enrichment, edge transparency indicates the number of shared genes between GO terms. The cyan group did not show any term enrichment. The position of the wild-type strain is marked with a black asterisk.

## Discussion

We have developed a highly efficient, low cost and rapid method, CRI-SPA, for parallelized transfer of a genetic feature from a haploid donor strain to any haploid yeast library strain. As an example, we have used CRI-SPA to transfer a five-genes cassette for betaxanthin production into nearly all strains of the YKO library. In fact, only 62 genes (1.3%) of the library consistently failed to produce any colonies through three screen repeats. As expected, these genes are associated with mating, cell division, and DNA repair, which represent functions that are essential for CRI- SPA (supplementary Table S3).

Gene transfer mediated by CRI-SPA is highly efficient and can be performed even without selecting for the genetic feature of interest. To this end, we stress that CRI-SPA mediated gene transfer is based on interchromosomal gene conversion. Hence, the donor sequence persists in the diploid state where gene transfer takes place allowing for a long tunable window of opportunity for transfer. This contrasts the scenario in classical gene targeting where the genetic feature is introduced via a linear DNA donor fragment or a self-cleaving plasmid system delivered by a SPA based method (Berry et al., 2020; Roy et al., 2018). In those cases, the time window where gene transfer can take place is limited to the lifespan of the donor fragment.

Importantly, CRI-SPA experiments are highly reproducible. Indeed, the *ade2 ade3* interaction was always scored when *ade2*Δ::hphNT1 was transferred to the appropriate subset of the YKO library. More impressively, when the YKO library-based betaxanthin strains generated in independent CRI-SPA screens were compared, we found that 86% and 75% of the top ∼200 hits generated in two trials using selection/selection and selection/no selection, respectively, were identical. Finally, we stress that hits identified in the screen are strongly enriched for GO-terms forming functional networks supporting a mechanistic consensus across hits. Hence, CRI-SPA is able to unravel novel physiology, which can be used in future metabolic engineering strategies for production of aromatic natural products like the morphine precursor L-DOPA.

Early in method development, we noted that CRI-SPA generates escapers, but at a low rate. Hence, in most cases, they will not contribute significantly to the phenotype of a colony. However, this is not the case if the escapers come with a fitness advantage where they may dominate at the end of the CRI-SPA procedure. Based on our analyses we suggest that most escapers are the result of incomplete gene transfer in cells that were aneuploid prior to CRI-SPA or by cells that underwent endoreduplication (Supplementary Figure S6). In agreement with this hypothesis, we essentially eliminated the background problem by adding a preincubation step on the sub-optimal carbon source raffinose prior to the SPA step. We speculate that this extra step extends the time-window for Cas9 induced gene conversion in a setting where the cell cycle is slowed down providing sufficient time to modify both copies of the target chromosome in these cells. We note that some mutants in the YKO collection may be prone to produce artifacts. For example, in the *ade2*Δ::hphNT1 transfer experiment, *scp160*Δ*::kanMX* was scored as a false positive. Interestingly, mutations in this gene are known to cause ploidy instability and to mate less efficiently (Gelin-Licht et al., 2012; Guo et al., 2003; Wintersberger et al., 1995) and we speculate that mutants with this phenotype may reduce the efficiency of CRI-SPA. In the absence of selection, additional background may be expected from two main sources. Firstly, unmated recipient cells passing through the CRI-SPA procedure would provide background and need to be eliminated. In our setup, this is done by selecting for a positive marker present on the CRI-SPA vector harbored by the CDS. Secondly, diploid cells that escape Cas9 induced gene conversion will result in background. It is therefore important that the Cas9-sgRNA nuclease efficiently cuts the target chromosome in all cells. Consequently, we recommend verifying Cas9-sgRNA cutting efficiency with TAPE (Vanegas et al., 2017). Moreover, the raffinose step is particularly recommended here to allow for prolonged Cas9-sgRNA nuclease activity in the diploid state.

Currently, a limitation of our method is that a recipient library needs to be *ura3* to be compatible with the CRI-SPA selection procedure. This could be bypassed by replacing the *Kl_URA3* markers in the donor strain by another counter-selectable marker like *amdS* (Solis-Escalante et al., 2013) to produce an even more flexible version of CRI-SPA.

Our successful marker-free gene-transfer experiment opens promising avenues for CRI-SPA. Bypassing the need for marker selection sets the stage for the transfer of point mutations as well as the multiplex transfer of several genetic features located at different positions in the genome at once. In another application, CRI-SPA could be used to transfer a genetic trait to libraries with different genetic backgrounds or even closely related species, as long as they can mate with the CDS. This is possible as CRI-SPA is independent of meiotic recombination and sporulation. To conclude, CRI-SPA is a powerful high-throughput data generation tool opening up a range of new exciting applications both in fundamental genetics and in genome engineering.

## Supporting information

Supp Table 3A Betaxanthin Top Hits

Supp Table 3B Betaxanthin Bottom Hits

Supp Table 3C Failing Genes

## Acknowledgements

The authors would like to thank Rodney Rothstein, Columbia University (NY, USA) for the strains W8164-2B and W8164-2C; and Irina Borodina, DTU (Denmark), for the plasmids pCfB2311 and pCfB2312. This project has received funding from the European Union’s Horizon 2020 research and innovation programme under the Marie Skłodowska-Curie grant agreement No 722287 (PAcMEN) to U.H.M, M.K.N, and J.M.D. The Danish National Advanced Technology Foundation (grant number 087-2012-3) to U.H.M.; the Villum Foundation and the Danish National Research Foundation (DNRF115) to M.L.; the Novo Nordisk Foundation Bioscience Ph.D. Programme grant No. NNF19SA0035438 to P.C. and the Fermentation-based Biomanufacturing Initiative, grant number NNF17SA0031362 supported N.S.

## Supplementary Material

### Strains and Media

All strains constructed in this work are listed in Supplementary Table 1. Strains W8164-2B and W8164-2C of the SPA method (Reid et al., 2008) were kindly provided by Rodney Rothstein (Department of Genetics and Development, Columbia University, USA). The YKO library was acquired from Invitrogen. YPD and synthetic complete (SC) media, and SC drop out media were prepared as described by), but with 60 mg/L L-leucine. For *URA3* counter selection SC plates were supplemented with 5-fluoroorotic acid (5-FOA) (1 g/L) and uracil (30 mg/L). To enhance the red phenotype of *ade2* cells, SC medium with only 4 mg/L adenine was used. To prevent *ade2* cells to be outcompeted by *ADE2* cells, growth media were supplemented with 40 mg/L adenine. Galactose and raffinose solutions were sterilized by filtration and used at 2% final concentration. For growth on solid medium, 20 g/L agar was added. For selection on solid medium, plates were supplemented with G418 (200 mg/L), nourseothricin (NTC) (100 mg/L) and hygromycin (hyg) (200 mg/L) as indicated. Half the concentration of the respective antibiotics were used in liquid cultivations. All strains were validated by diagnostic colony PCR to ensure correct integration at the intended chromosomal locus. For an experimental overview of CDS construction, see Supplementary Figure S1.

### Plasmid construction and PCR

All cloning was done by USER-cloning (Geu-Flores et al., 2007; Nour-Eldin et al., 2006) employing uracil-specific excision reagent (USER™) enzyme from New England Biolabs. DNA polymerases were purchased from Thermo Scientific and used according to the supplier’s instructions. Amplification of DNA for USER cloning was carried out using Phusion U Hot Start DNA Polymerase. Standard PCR amplifications for cloning were run using Phusion Hot Start II High-Fidelity DNA Polymerase. In the case of colony PCR, Taq DNA Polymerase was used. Purification of DNA fragments obtained by PCR or from agarose gel bands was done using the illustra GFX PCR DNA and Gel Band Purification Kit (GE Lifesciences).

*cas9* vector: pHO8 contains a *cas9* gene targeting substrate for integration into chromosomal integration site X-3 (Mikkelsen et al., 2012). pHO8 was made by amplifying *cas9* from plasmid pCfB2312 (Stovicek et al., 2015) with primers HOP89 and HOP90. The fragment was then inserted into a linear vector fragment of the integrative plasmid pCfB257 (Jensen et al., 2014) obtained by PCR using primers HOP91 and HOP92.

### Betaxanthin vectors

The integrative plasmids for inserting betaxanthin-synthesizing genes in the XII-5 site were constructed using the Ant_E113 plasmid as template (Coumou et al., 2017). pBTX1 was built to exclude ARO4K229L and ARO7G141S and was obtained by PCR with primers HOP258 and HOP259. This linearised plasmid was closed by USER assembly. pBTX2 was built to include ARO4K229L and ARO7G141S and was obtained by PCR with primers PR_DIV2288 and ANT_P494 and ligated into pCfB2909 with USER assembly.

### sgRNA coding Vectors

For construction of the CRI-SPA vector pHO-ADE2 targeting the *ADE2* locus, the target sequence in pCfB3050 (Jessop-Fabre et al., 2016) was excluded by linearizing the plasmid with uracil-containing primers HOP60 and HOP61 and replaced by an insert containing the 20 bp *ADE2* targeting sequence. The insert was obtained by annealing two single stranded oligonucleotides HOP190 and HOP191. For insertion of an additional *KlURA3* marker between *ADE2* and the telomere, an sgRNA plasmid for targeting an intergenic sequence downstream of the ADE2 locus (pHO22) was constructed as described above but instead annealing the two oligonucleotides HOP173 and HOP174. Similarly, the sgRNA plasmid for targeting the intergenic sequence downstream the XII-5 integration site (pHO25), was produced by inserting a 20 bp targeting sequence obtained by annealing the two oligonucleotides HOP183 and HOP184. pHO-XII-5, expressing an sgRNA targeting XII-5, was made by exchanging by USER cloning the NatMX marker cassette of plasmid pCfB3050 (Jessop-Fabre et al., 2016) for a hphNT1 cassette PCR amplified from pMEL12 (Mans et al., 2015) using uracil-containing primers HOP217 and HOP218.

### Construction of Yeast Strains

All yeast transformations were carried out using the LiAc / SS Carrier DNA / PEG method described by Gietz and Schiestl (Gietz and Schiestl, 2007). When antibiotic resistance genes were used as markers, cells were allowed to recover for 2 hours in liquid YPD prior to plating on solid selective media.

Type 1 Universal CRI-SPA strains, Type 1 UCSs, were made by inserting *cas9* into chromosomal integration site X-3 of W8164-2B (*MATα*) and W8164-2C (*MAT<u>a</u>*) using a gene targeting substrate liberated from pHO8 by digestion with NotI (Thermo Scientific). UCS type 2 strains were made from UCS type 1 strains by inserting an additional copy *of KlURA3* between expression site XII-5 and the telomere. Insertion was achieved via co-transforation with sgRNA plasmid pHO25 and a linear KlURA3 repair substrate liberated from pCfB390 (Jensen et al. 2014) using primers HOP181 and HOP182. CRI-SPA donor strains were made from UCS strains. CDS-*ade2*Δ was constructed by first transforming UCS type 1 with sgRNA plasmid pCfB2311 (Stovicek et al., 2015) and a linear repair substrate for disruption of *ADE2* via insertion of the *hphNT1* cassette, which was generated by PCR using plasmid pMEL12 and primers HOP64 and HOP65. The resulting strain was then further modified to harbor an additional *KlURA3* marker between the disrupted *ADE2* and the telomere by transformation with sgRNA plasmid pHO22 and a linear *KlURA3* repair substrate amplified from pCfB390 (Jensen et al 2014) by primers HOP175 and HOP176, yielding CDS-*ade2*Δ .

CDS-btx1 was constructed from the UCS type 2 strain by transforming with the integrative plasmid pBTX1 harboring both the betaxanthin-synthesizing genes CYP76AD1 and DOD, and an sgRNA expression cassette for targeting the XII-5 site. CDS-btx2 was constructed by co-transforming the UCS type 2 strain with the linearised plasmid pBTX2 and pHO-XII-5.

### CRI-SPA high-throughput pin replication protocol

Automatic pin replication was carried out using the high-throughput pinning robot ROTOR HDA from Singer Instruments (United Kingdom), along with replica pinning pads (RePads) and rectangular petri dishes (PlusPlates) from the same company.

For parallel mating of the Universal Donor Strain to the strains of the yeast deletion library, a single colony of the UDS was inoculated in 25 mL YPD supplemented with the appropriate antibiotic in order to maintain selective pressure for the gRNA plasmid. The UDS was grown overnight and washed by centrifugation and resuspended with 25mL sterile water to remove the antibiotic. 150uL of cell suspension was dispensed in wells of a 96 well plate serving as a source plate for the screen. From this source plate, the UDS was pinned in 1536 format on the mating plates. The arrayed strains of the deletion collection were pinned on top of the UDS. Plate shuffling was introduced at this stage to address experimental artifacts (see below). Mating was allowed to proceed from 8 hours to overnight. Strains were then pin replicated onto a series of selective plates as outlined in Figure 1.

### Image processing, normalization and filtering

Several experimental factors (agar, plate, neighbor and edge effects) affect colony metrics in high throughput array screens. These need to be corrected so that any two colonies become directly comparable regardless of their positions within the screen. Instead of placing the four 384 arrays side by side on the same plate, each of the 4 arrays was randomly assigned a different plate within the screen Fig. S6). In this setting, quadruplicates do not sit adjacent to each other and are exposed to a much greater diversity of neighbors (up to 13*9) than in the unshuffled mode (9, including themselves). In addition, the 4 quadruplicates grow on different plates, removing the agar gradient effect from our signal. Importantly, the row-column positions of the quadruplicates are preserved in this setting (Supplementary Fig. S6)

This setup has the advantage of randomizing the effects of agar and neighbors without the need of further statistical processing. The edge effect, the strongest artifact, is however not randomized. This would greatly increase the within-quadruplicate variation and is undesirable. Rather, this effect was corrected in the 5 outer frames with a simple dot product as previously described (Collins et al., 2006).

Next, we built a bespoke image analysis pipeline extracting colony size and color and correcting for those external factors. Colony size and color were extracted with functions adapted from Pyphe (Kamrad et al., 2020; scripts can be found at https://github.com/pc2912/CRI-SPA_repo). To quantify yellow color intensity, the image was converted to HSV and yellow intensity was defined as the geometric mean of Saturation and Value (Fig. S6). This assumed that the colony’s Hue was fixed (i.e. colony can only harbour variants of yellow) which was our observation on YPD media.

Outliers within quadruplicates were detected with Grubbs’ test for outliers and deleted. Genes represented by two colonies or less were deleted from the analysis.

### Gene Enrichment Analysis and GO term graphs

Gene enrichment analysis was performed using GOATOOLS (Klopfenstein et al., 2018) for gene groups as indicated on Figure 5. The default Benjamini-Hochberg correction was used to account for the number of tests, and a P-value of 0.05 was used as a threshold to accept GO terms as enriched.

For visualizing results at a systems-level, a graph was constructed where nodes representing GO terms were connected if they shared associated genes. The size of the nodes was drawn as a function of the significance of the term enrichment and the edge opacity was drawn in relation to the number of genes shared between terms. Nodes were colored according to their description or the description of their ancestors (i.e. descriptions contained key words such as ‘mitochondria’, ’translation’, ect.). Finally, the resulting networks were built in NetworkX, transferred to Cytoscape (Shannon et al., 2003) with py4cytoscape and minor manual adjustments were made for visibility (scripts available at https://github.com/pc2912/CRI-SPA_repo)

## SUPPLEMENTARY FIGURES

**Supplementary Figure S1.**
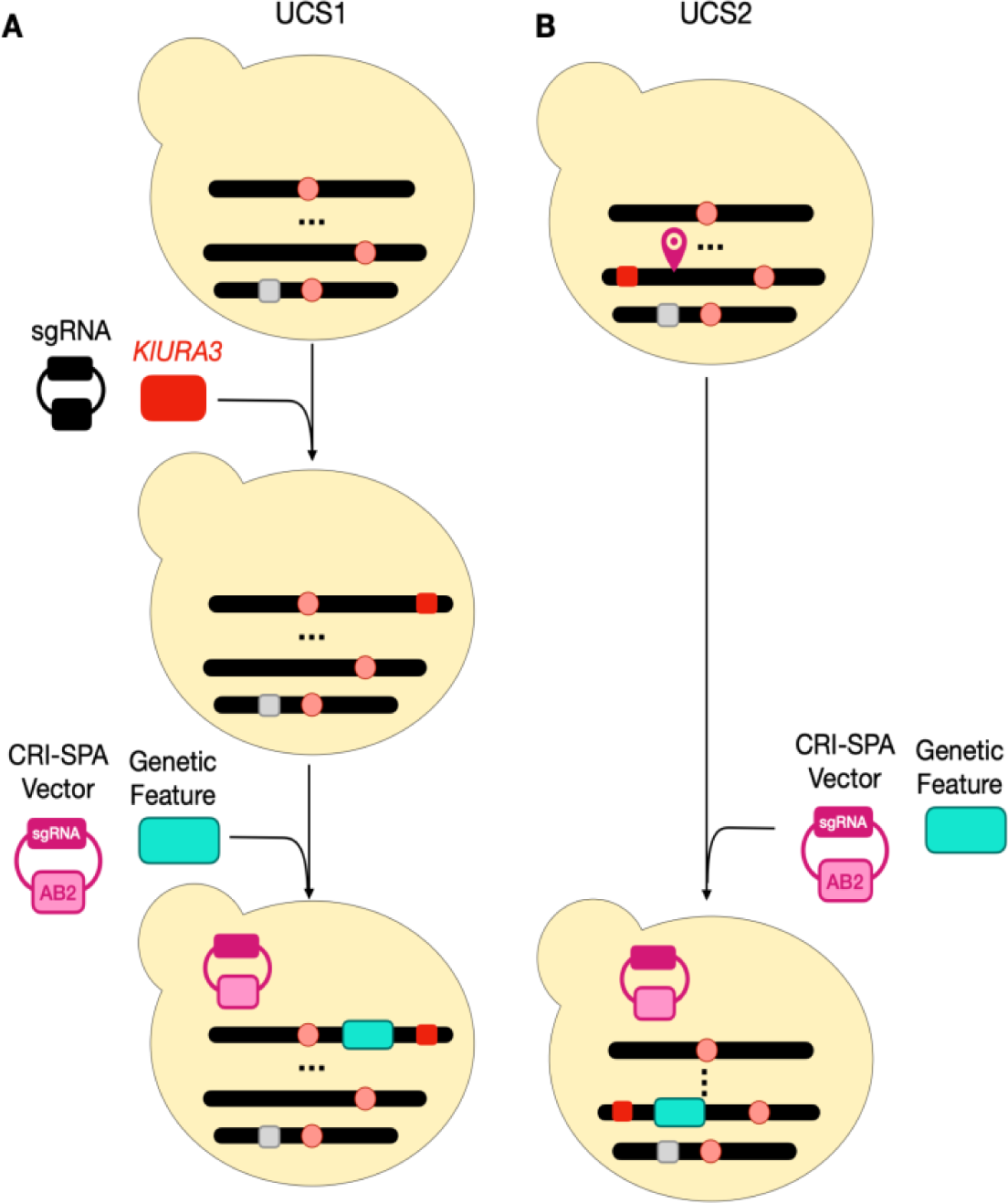
CDS construction. Construction of a CDS depends on the nature of the genetic feature of interest. A) If the feature is to be transferred from a user-defined site, the CDS needs to be constructed in two steps using a UCS type 1 strain (UCS1). Firstly, a URA3 marker needs to be incorporated on the telomeric side of the transfer site with the help of a sgRNA plasmid. After curing of the sgRNA plasmid, the genetic feature is transformed together with the CRI-SPA vector. B) If the genetic feature of interest is a gene or an entire genetic pathway that needs to be expressed from a defined expression site, the CDS can be constructed in a single step using UCS2. UCS2 comes with a pre-integrated KlURA3 marker between the defined expression site XII-5 and the corresponding telomere. Moreover, an efficient CRI-SPA vector for this site is also available (pHO-XII-5 see Table S2). For convenience, UCS1 and UCS2 strains are available in both mating types (see Table S1)

**Figure S2:**
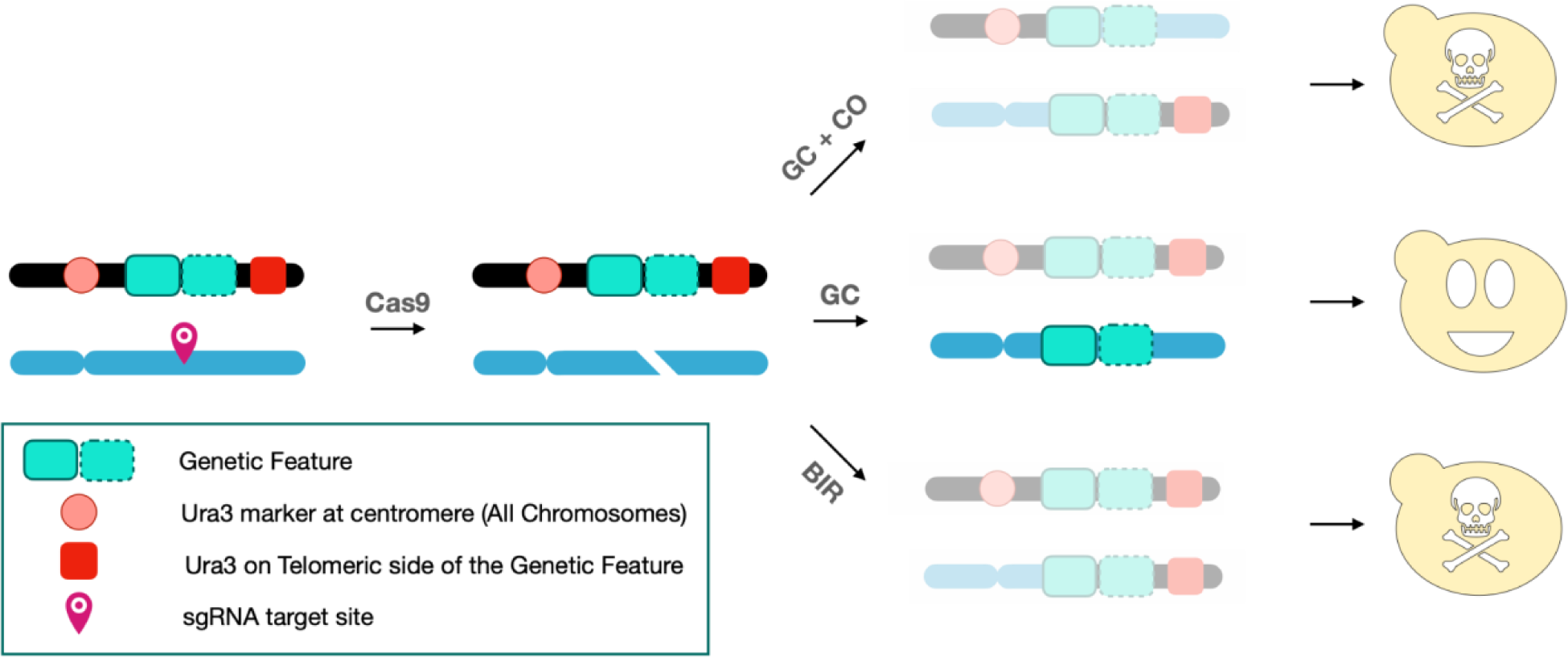
CAs9 induced Crossover and break induced replication produce chimeric strains. In CRI-SPA, transfer of the desirable genetic information from the donor to the target site by homology directed repair of a DNA DSB in the target site is performed by gene conversion (GC). However, homology directed repair of the DNA DSB may also occur via GC accompanied by a crossing over event (GC + CO) or by break induced replication (BIR). In both cases, the final strain after CRI-SPA will contain an undesirable chimeric chromosome consisting of donor and target chromosome sections. In our CRI-SPA system, strains created by GC + CO or BIR are counterselected in step 4 of the CRI-SPA procedure (5-FOA counter selection) as we have incorporated a *K. lactis* URA3 marker between the donor site and the telomere; and this *URA3* marker will be transferred to the target chromosome in case that repair of the CRISPR induced DNA DSB involves GC + CO or BIR.

**Figure S3.**
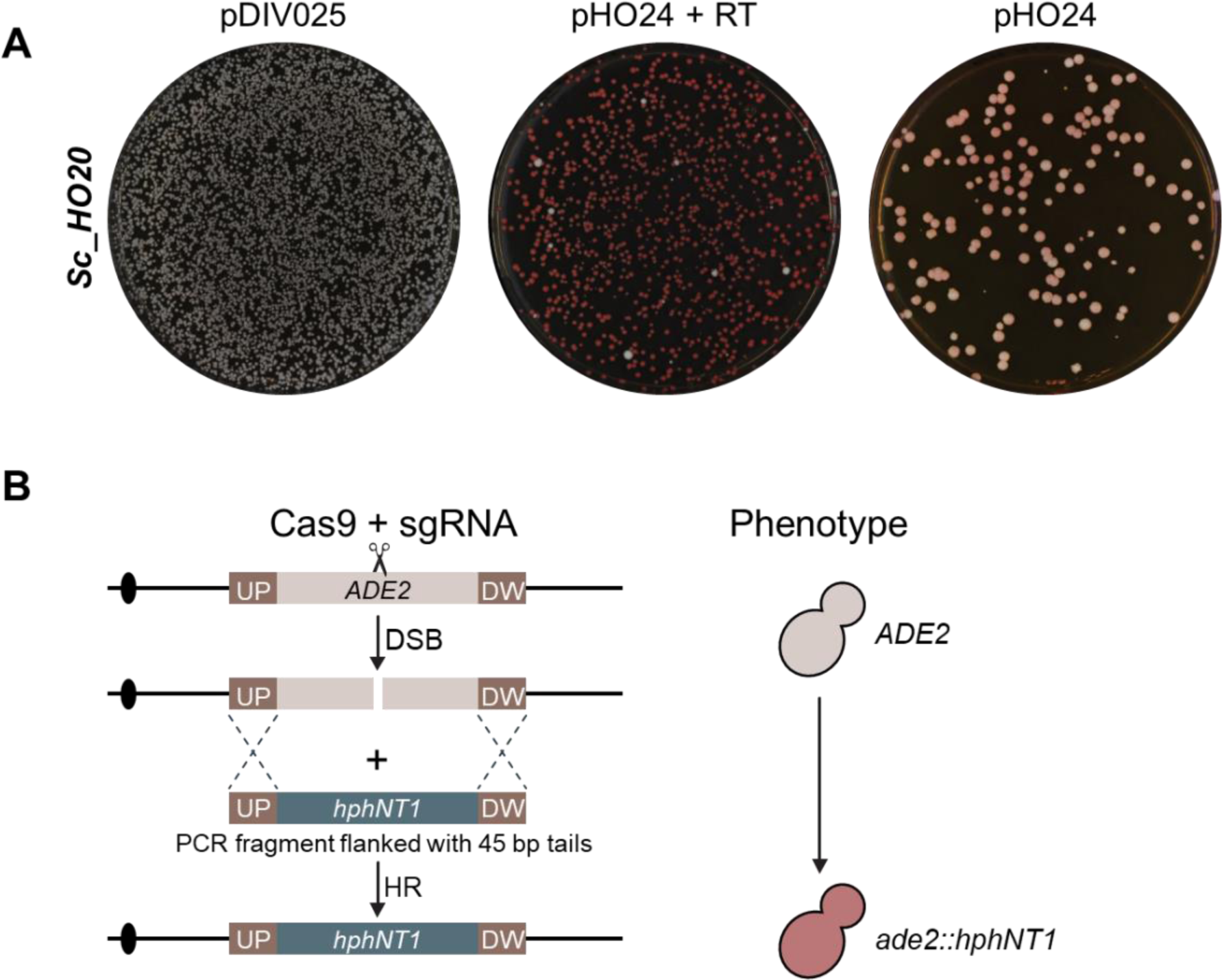
TAPE experiment to assess Cas9/gRNA cleavage efficiency of a target protospacer. A TAPE (tool <u>a</u>ssess <u>p</u>rotospacer <u>e</u>fficiency) experiment (Vanegas et al., 2017) was performed to determine how efficiently the CRISPR nuclease Cas9/*ADE2-*gRNA cuts the *ADE2* locus in a UCS1 strain (UCS1α). Panel **A** shows the numbers of transformants obtained after transformation of UCS1α with an empty plasmid pDIV025, and with pHO24 encoding the *ADE2*-sgRNA, as well as after co-transformation with pHO24 and a *ade2Δ::hphNT1* PCR fragment flanked with 45 bp long UP and DW targeting regions (Panel **B**) that serves as repair template (RT) for breaks induced at *ADE2*. The amounts of vector DNA used in the three transformation experiments were stoichiometrically the same. The high number of transformants obtained with pDIV025 relative to the number obtained with pHO24 indicates that lethal DNA DSBs are produced by the Cas9/*ADE2-s*gRNA CRISPR nuclease in UCS1α transformed with pHO24. This conclusion was supported by two observations. Firsty, the number of transformants obtained after transformation with pHO24 increases to match the number of transformants obtained with pDIV25 when the repair fragment was included in the reaction allowing for efficient DNA DSB repair by homologous recombination. Secondly, virtually all transformations obtained in the co-transformation experiment were red as *ADE2* was deleted.

**Figure S4.**
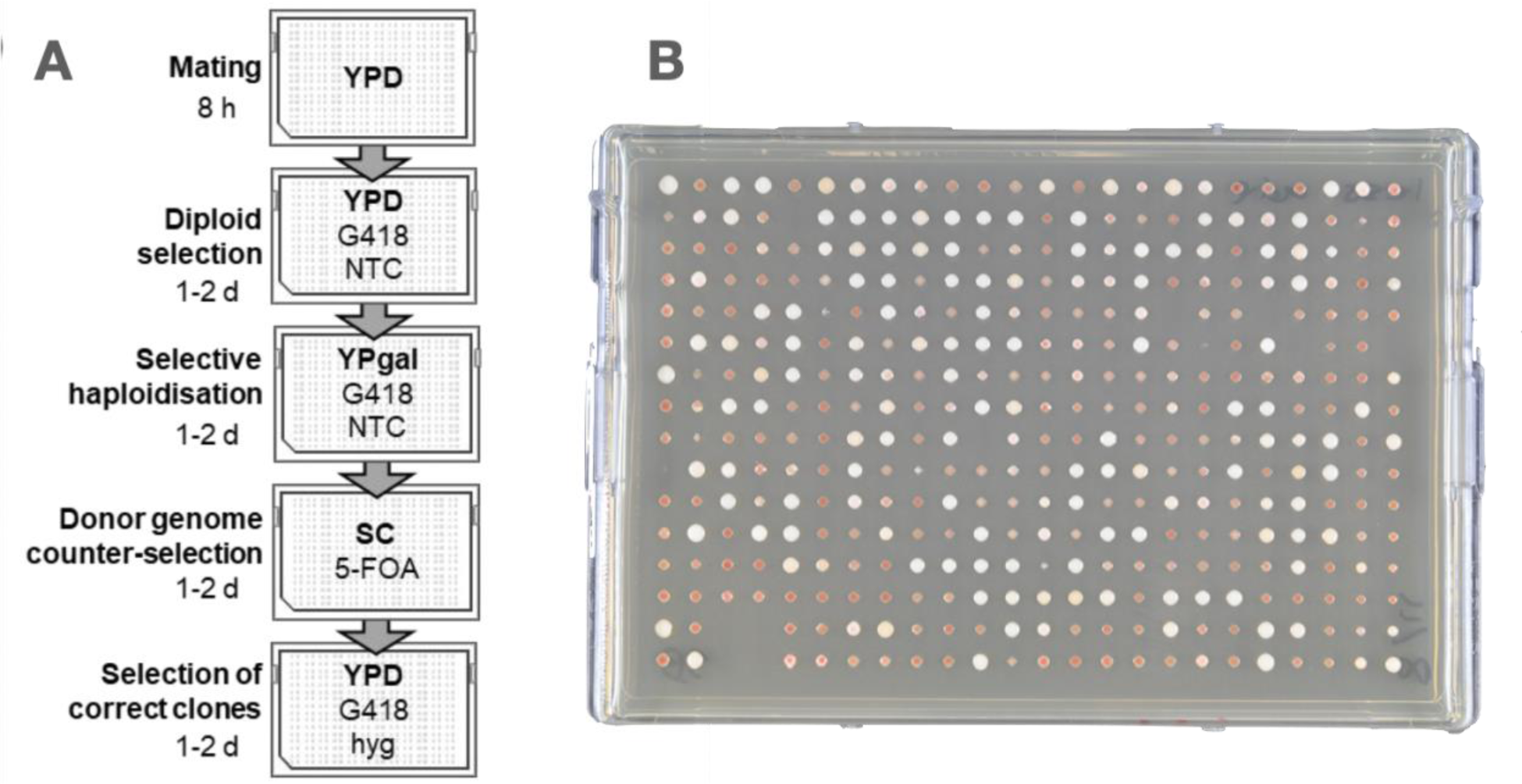
Initial test of a CRI-SPA plating protocol for deletion of the ADE2 gene in a subset of the yeast deletion collection. **A)** Initial replica plating procedure, adapted from Reid et al., 2011, with an additional step to select for diploids. For mating, a subset of the yeast deletion library (plate 9, 384 format) pinned onto a lawn of CDS-*ade*2Δ. **B)** Plate produced by the procedure.

**Figure S5:**
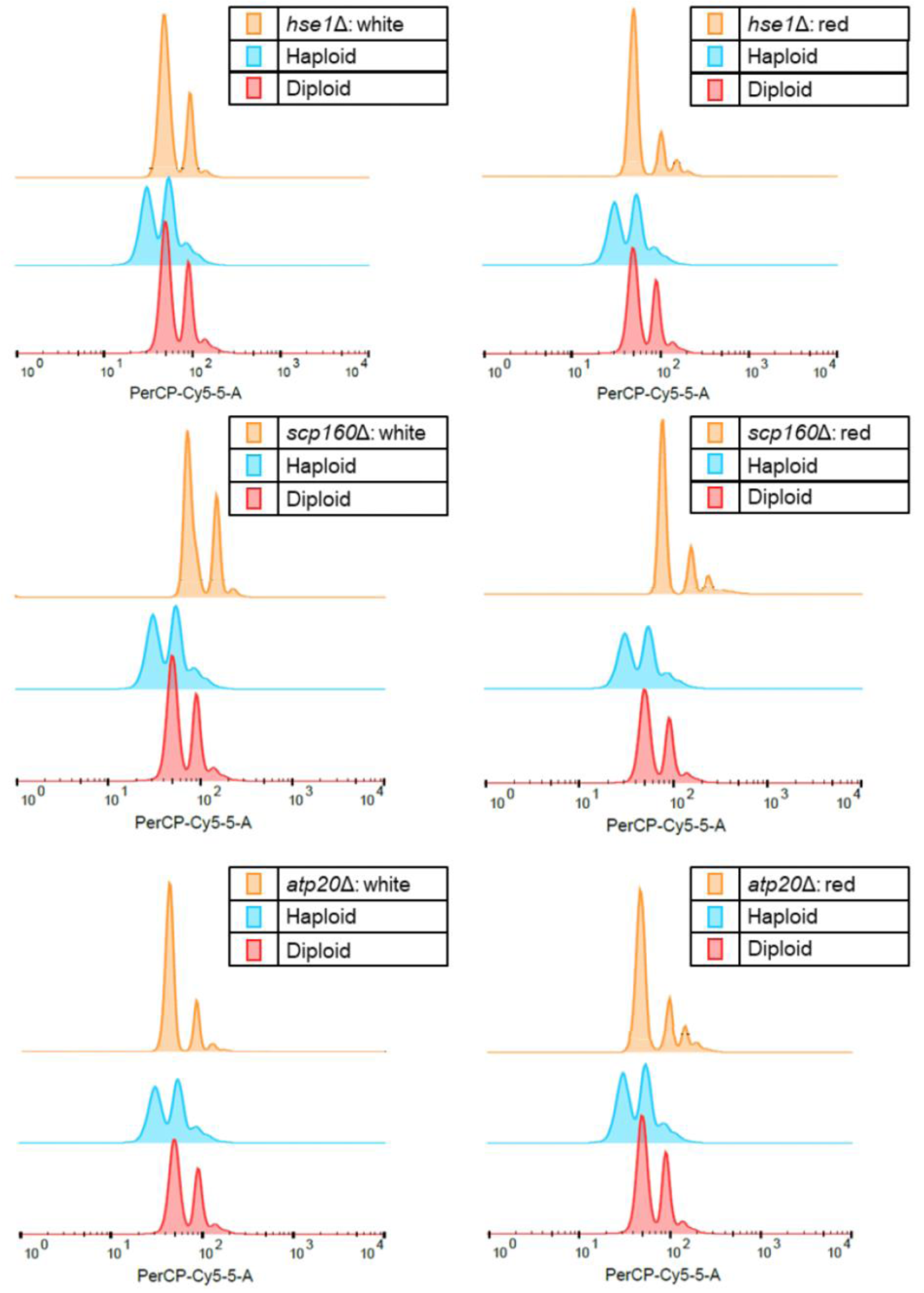
flow cytometry for ploidy. Strains originate from the plating procedure in Figure S4 and in each case, a white and a red single colony of the respective strain was analyzed. Cells were stained with propidium iodide and the histograms show the distribution of cells in accordance to their relative DNA content when analyzed by flow cytometry. Ploidy was assessed by comparing histograms of each clone to that of a known haploid and diploid reference strain.

**Figure S6.**
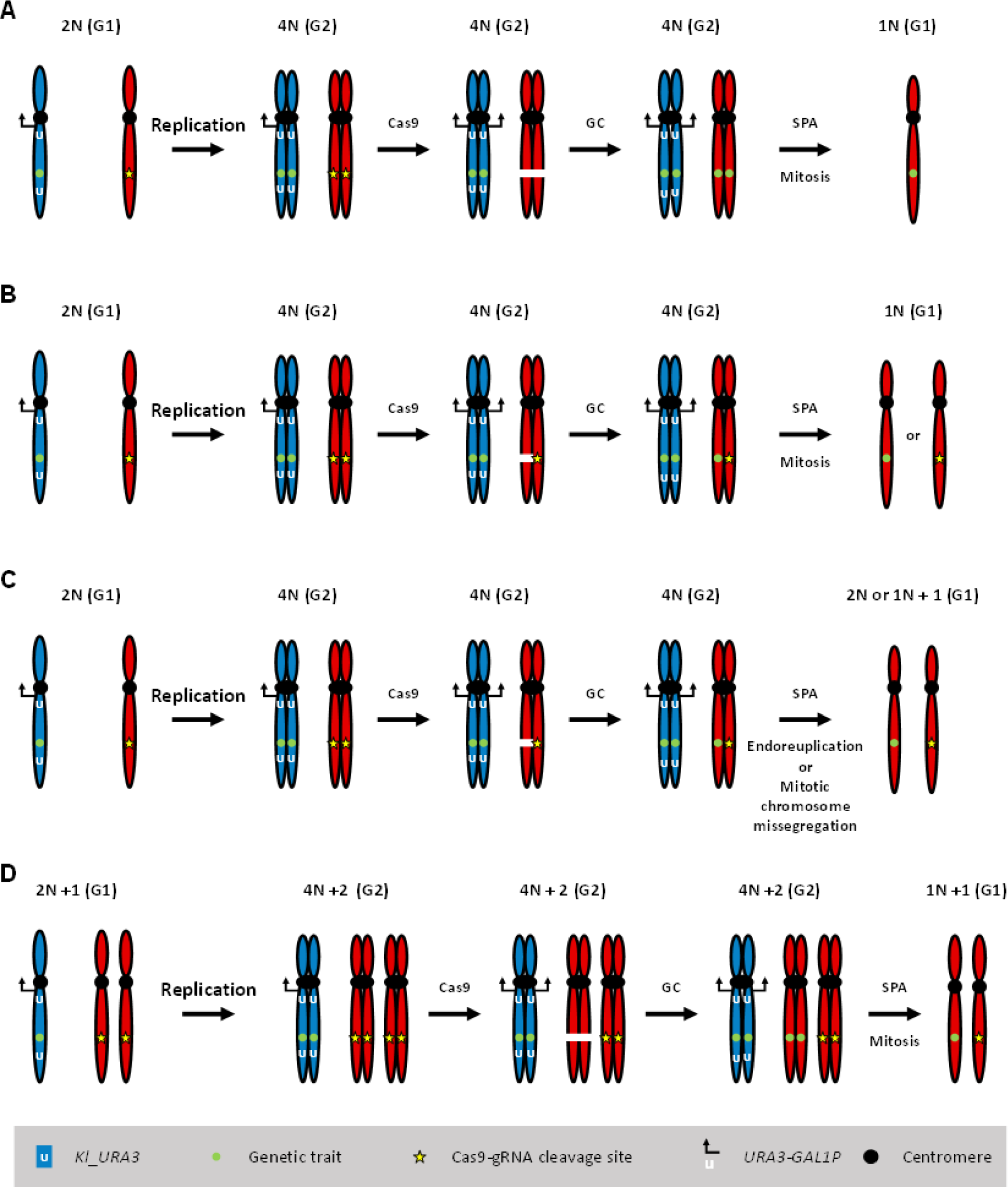
Models explaining background cells generated by CRI-SPA. Background may be produced in cells containing an extra copy of the target recipient chromosome. The extra chromosome may be due to the recipient strain being aneuploid from the beginning, or due to endoreduplication during SPA. If Cas9 only cleaves the target sequence in one of the two target chromosomes during CRI-SPA, CRI-SPA mediated transfer of a genetic trait from a donor chromosome (blue) to the corresponding recipient chromosome (red) is incomplete. Different scenarious are illustrated in the panel below with the diploid strain formed by mating of the CDS to a recipient strain during CRI-SPA as a starting point. **A)** in normal CRI-SPA, Cas9 will cleave both recipient target chromatids in the diploid cells formed by fusion of donor and recipient cells. Repair of the breaks by gene conversion (GC) transfers the genetic trait from a donor chromatid to the corresponding recipient chromatids. Haploidization by SPA generates recipient chromosomes Containing the genetic trait of interest. **B)** If CRISPR mediated GC takes place after the target locus has been replicated, the possibility exists that only one of the two recipient chromatids may be cleaved by Cas9 in the timeframe of the CRI-SPA process where Cas9 is active. If so, only one of the two recipient chromatids will receive genetic information from the corresponding donor chromatid. In this case cells may contain either a recipient chromosome containing the desired genetic trait, or one that does not. If the genetic trait is selected for, the latter type of cells will die. If the genetic trait is not selected for, the latter type of cells will survive and produce background. **C)** Same scenario as in B) except that the process does not continue with a perfect mitosis, rather the entire genome or the target chromosome is duplicated as the result of endoreduplication or the number of target chromosomes is doubled due to chromosome missegregation. As a result, cells containing heterozygous recipient disomes are formed where only one chromosome contains the desired genetic trait. Such cells are expected to survive in the presence or absence of selection for the genetic trait to produce background. **D)**. The starting recipient cell is aneuploid and contains an additional target chromosome. Like in the other scenarios, incomplete cleavage by Cas9 produce one modified and one un-modified target chromosome in the final cell.

**Figure S7.**
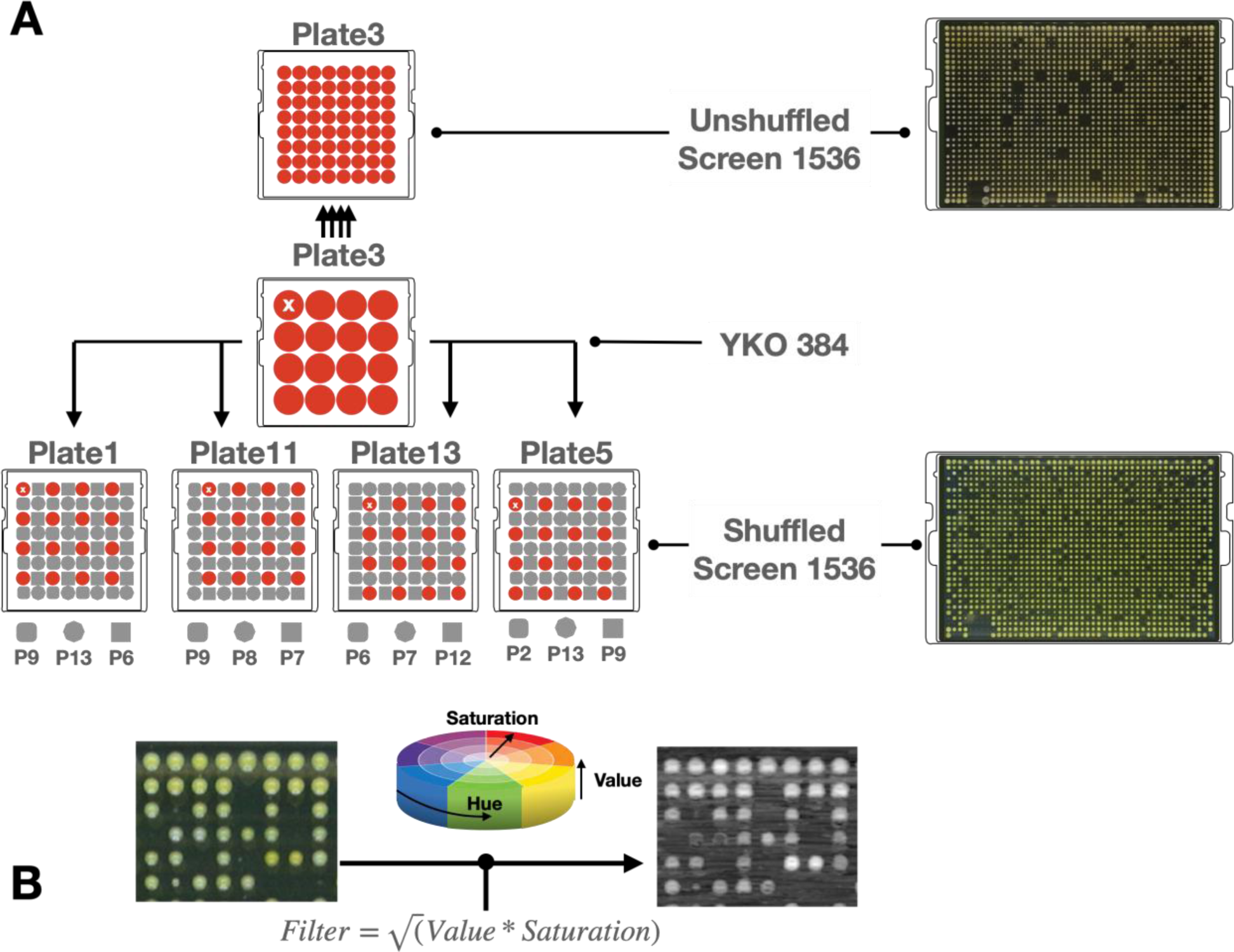
Colony Shuffling Scheme and Color Filtering. A) Left: first version of the screen. During expansion from 386 to 1544 format, individual mutant colonies are pinned as colony quadruplicates in a square of the same plate. As a result, each quadruplicate colony is affected by identical agar quality and similar neighbor effects. Blue square shows YOR1 mutant. Right, second version of the screen. During expansion from 386 to 1544 format, individual mutant colonies are shuffled across random screen-plates. For example, YKO plate 1 is pinned on screen plates 7, 3, 6 and 10. This increases the diversity of neighbors and reduces potential agar gradient biases. B) Heuristic to quantify yellow intensity. Yellowness was quantified as the geometric mean between Value and Saturation in the HSV domain.

**Figure S8.**
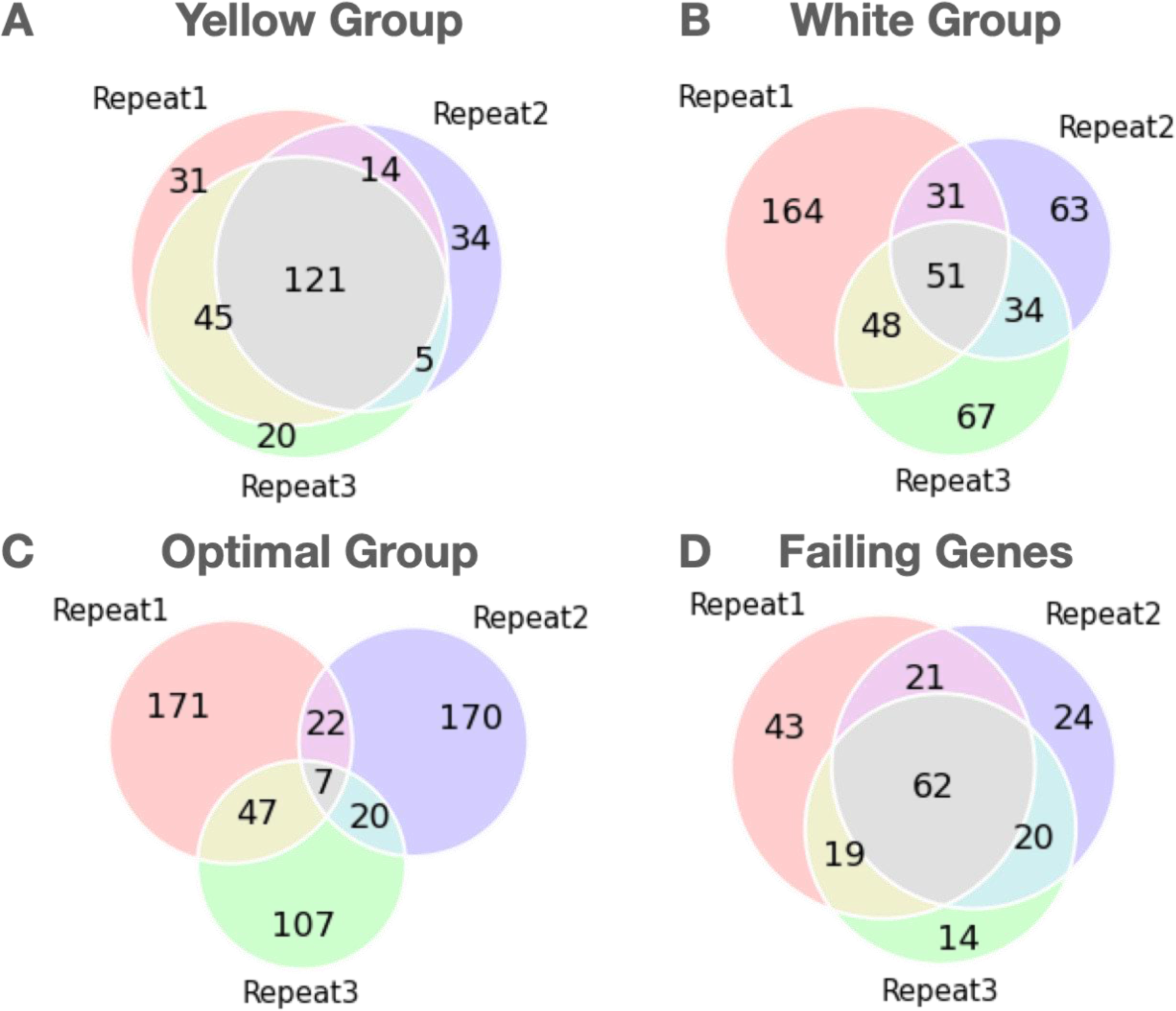
Screen repeats overlap. Overlap in gene groups between three screen repeats, A) Yellow Group, top betaxanthin (Yellow intensity > 1.2 std above screen mean). B) White Group, lowest betaxanthin (Yellow intensity < 1.2 std below screen mean). C) Optimal group (0.8 < Yellow intensity < 1.2 std above screen mean). D) Failing genes, genes producing 0 colonies after CRI-SPA procedure.

**Figure S9.**
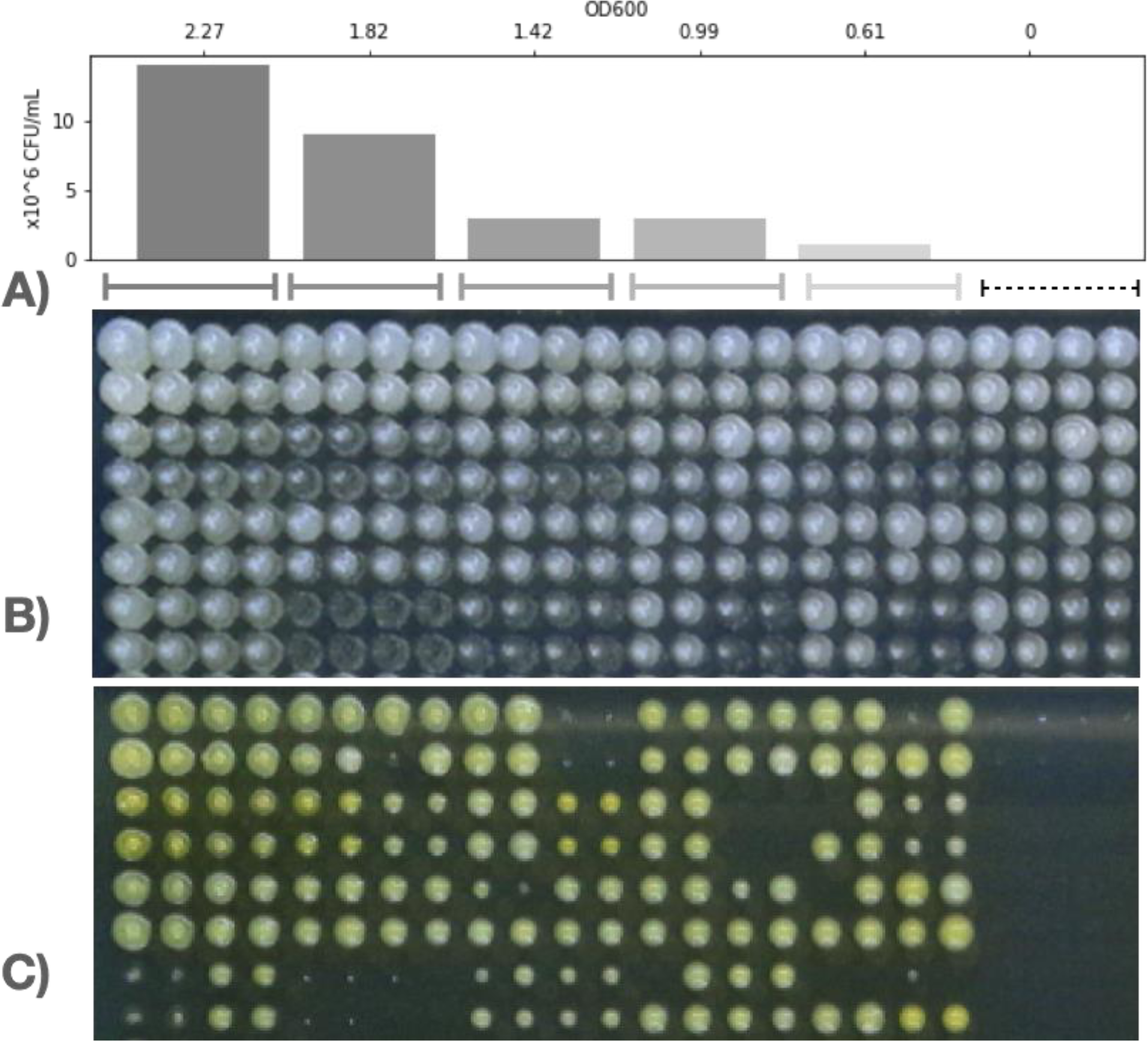
Pilot screen for optimal starting UDS concentration. The UDS was pinned for mating from a 96well plate at different ODs. A) Starting OD of the suspension corresponding to the columns of 4 colonies on the underlying pictures. B) Plate picture during mating step. Note, the colonies are white because the YKO library is more abundant than the UDS at this step. C) Final CRI-SPA plate: CRI-SPA succeeded across the range of OD tested. Empty spots correspond mostly to colonies where the YKO was absent at the mating step.

**Table S1:**
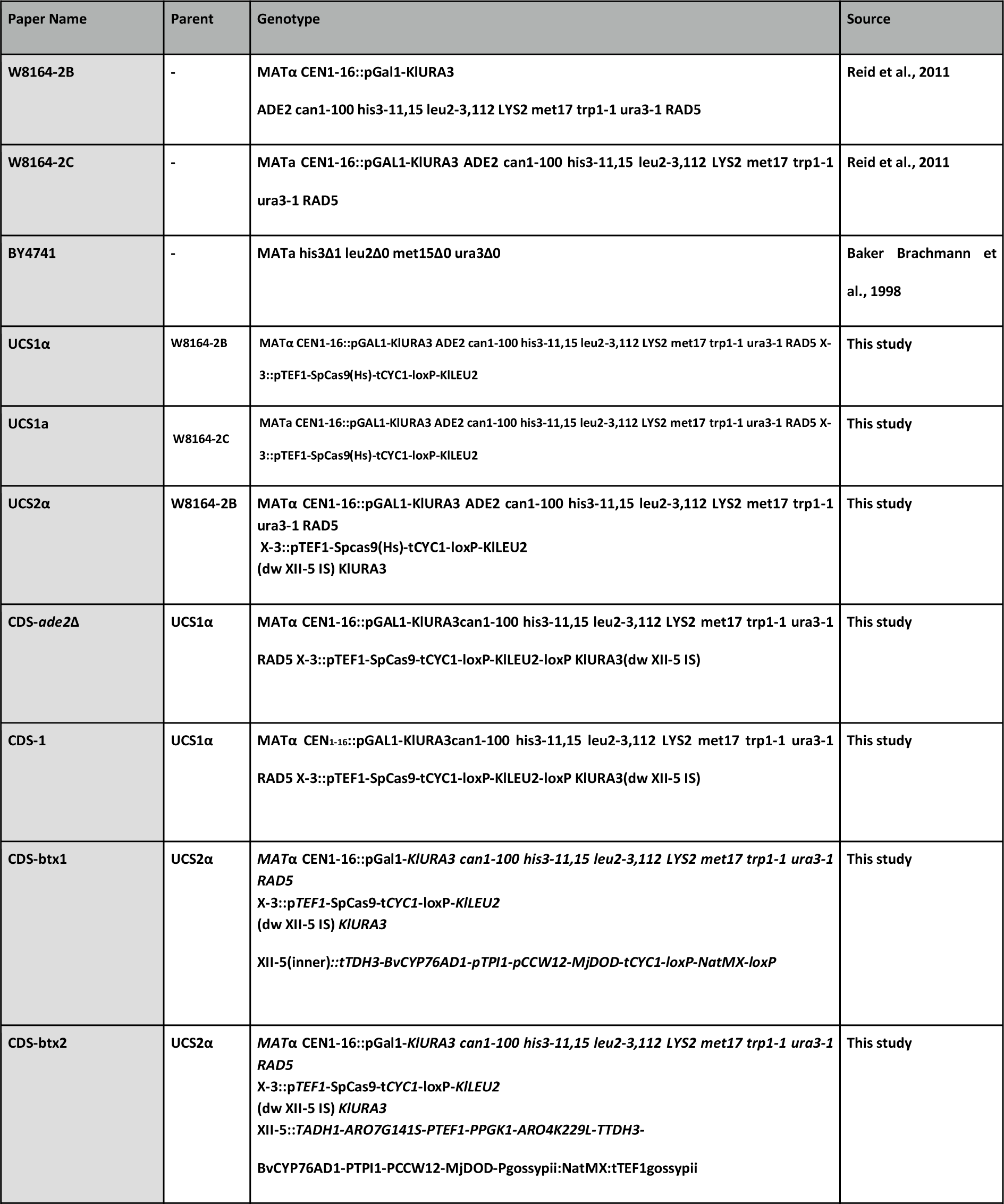
Strains used in this study.

**Table S2A:** Vectors used in this study.

| Plasmid Name | Yeast Marker | Type | Genetic Element |
| --- | --- | --- | --- |
| pHO-XII-5 | hphNT1 | 2μ | pSNR52-sgRNA(XII-5)-tSUP4 |
| pHO-ADE2 | hphNT1 | 2μ | pSNR52-sgRNA(ADE2)-tSUP4, NatMX |
| pHO8 | – | Integrative, X-3 | pTEF1-SpCas9-tCYC1 |
| pBTX1 | NatMX, URA3 | Integrative, XII-5 | pSNR52-sgRNA(XII-5)-tSUP4, tTDH3-BvCYP76AD1-pTPI1-pCCW12-MjDOD-tCYC1 pTEFagossypii::NatMX::tTEFagossypi |
| pBTX2 | NatMX, URA3 | Integrative, XII-5 | TSUP4ARO7G141S-PTEF1-PPGK1-ARO4K229L-TTDH3-BvCYP76AD1-PTPI1-PCCW12-MjDOD pTEFagossypii::NatMX::tTEFagossypi |
| pHO25 | NatMX, URA3 | 2μ | pSNR52-sgRNA-tSUP4 |

**Table S2B:**
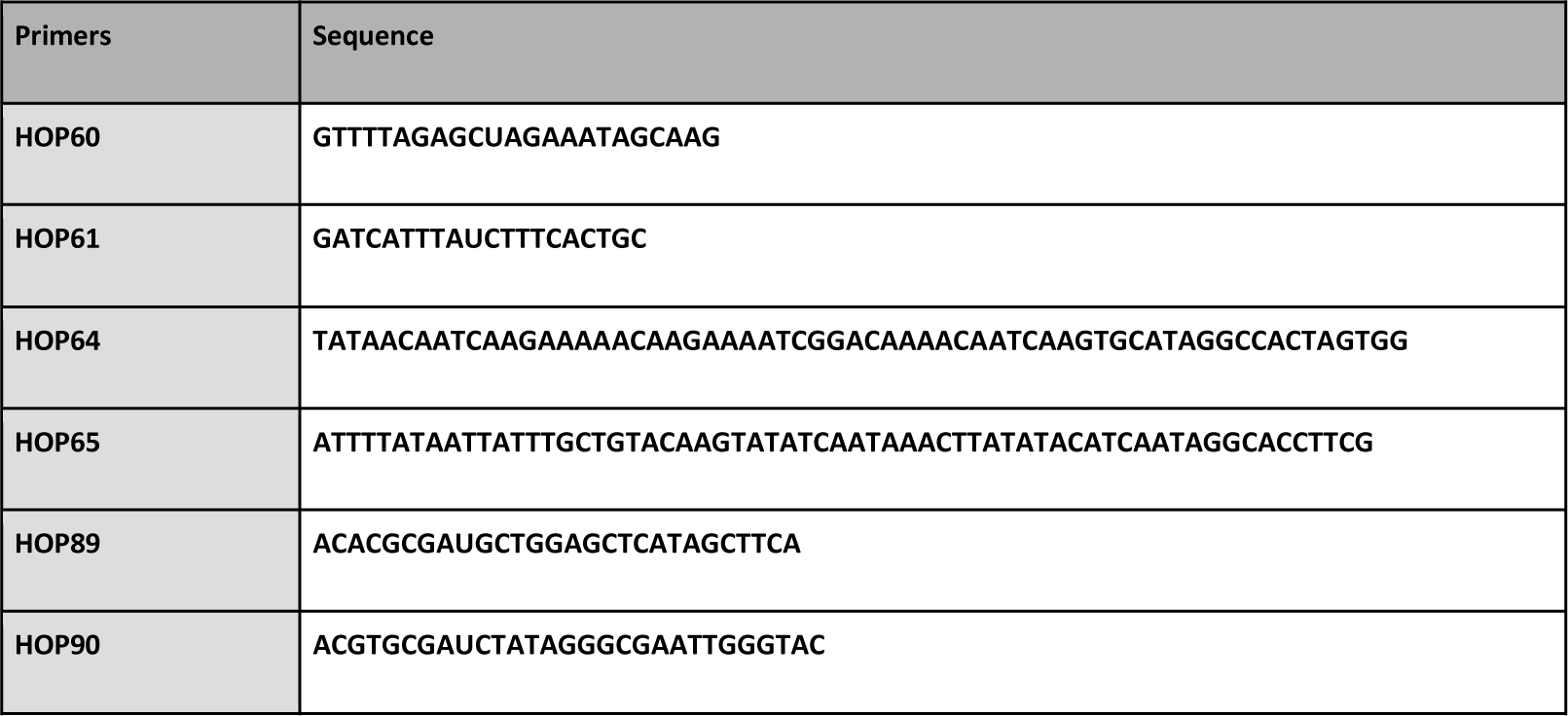
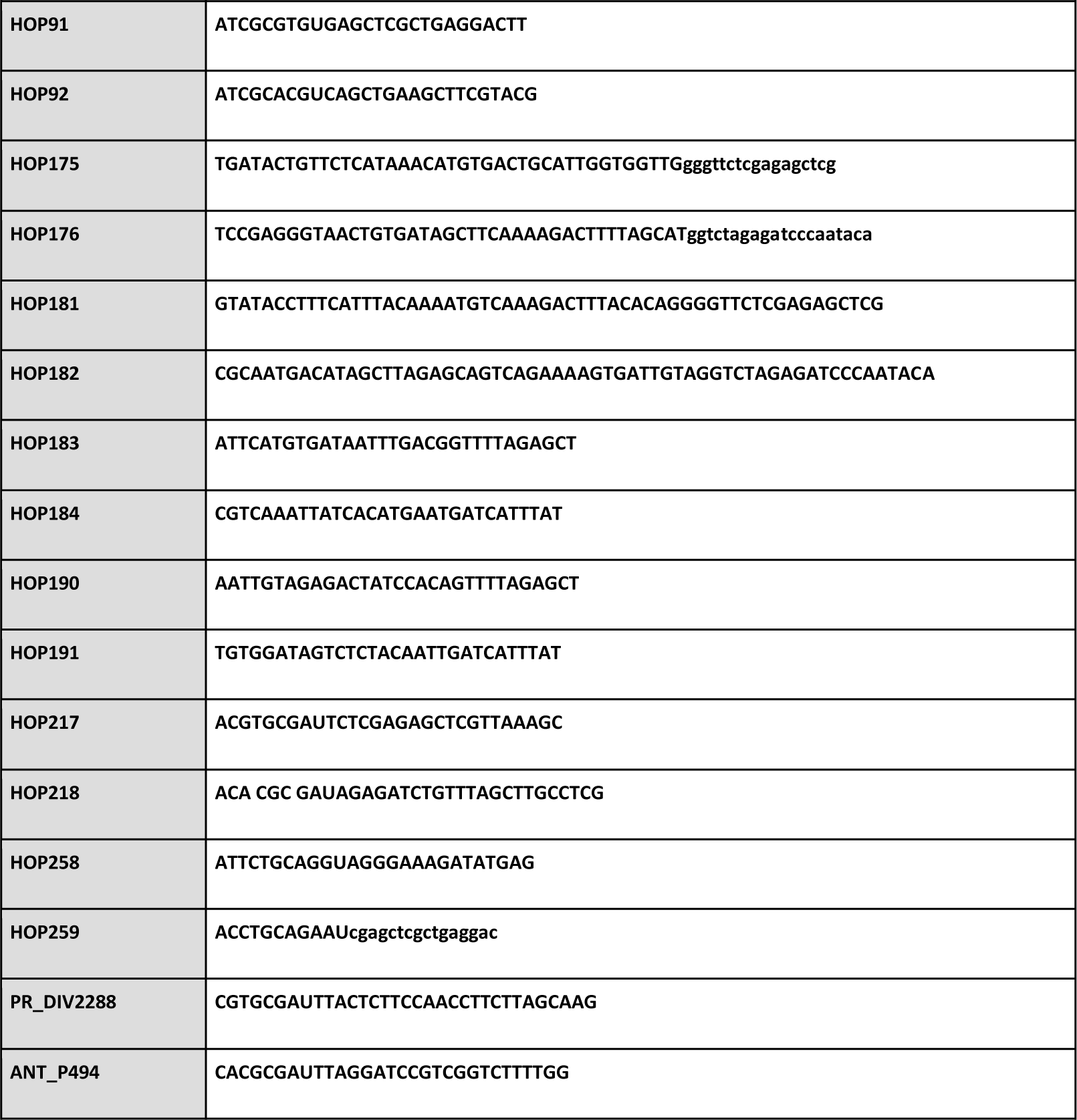
Vectors used in this study.

## Notes

### Competing Interest Statement

The authors have declared no competing interest.

### Summary of Updates

BioRxive author order was corrected. U. Mortensen, the corresponding author was placed last.

